# An nf-core framework for the systematic comparison of alternative modeling tools: the multiple sequence alignment case study

**DOI:** 10.1101/2025.03.14.642603

**Authors:** Luisa Santus, Jose Espinosa-Carrasco, Leon Rauschning, Júlia Mir-Pedrol, Igor Trujnara, Alessio Vignoli, Leila Mansouri, Athanasios Baltzis, Evan W. Floden, Paolo Di Tommaso, Edgar Garriga, Adam Gudyś, Sebastian Deorowicz, Cameron Gilchrist, Martin Steinegger, nf-core community, Cedric Notredame

## Abstract

The computational complexity of many key bioinformatics problems has resulted in numerous alternative heuristic solutions, where no single approach consistently outperforms all others. This creates difficulties for users trying to identify the most suitable tool for their dataset and for developers managing and evaluating alternative methods. As data volumes grow, deploying these methods becomes increasingly difficult, highlighting the need for standardized frameworks for seamless tool deployment and comparison in HPC environments. Multiple sequence aligners (MSAs) rank among the most commonly employed modeling techniques in bioinformatics, playing a crucial role in applications such as protein structure prediction, phylogenetic reconstruction, and variant effect prediction. The NP-hardness of MSAs makes them a major example of problems where heuristics stand central, as no optimal solution can be currently obtained, within the limits of operational computational requirements. Here, we present a pilot design of an nf-core framework for streamlined tool deployment and rigorous performance evaluation focusing on the MSAs software ecosystem. By integrating the most popular MSA tools and focusing on a modular, and extensible architecture, we aspire to provide a key platform supporting MSA deployment, evaluation, and algorithmics development to the MSA community, and a proof-of-principle to the wider bioinformatics community.

## Introduction

The massive generation of biological data over the last decades has resulted in an increasing reliance on fast, approximate heuristic algorithms in bioinformatics tools. Multiple sequence alignment (MSA) is an example of a problem where alternative heuristic methods are available. Yet, this methodological configuration is far from being an isolated case. Indeed, data growth has been such that even for tasks having exact solutions, such as database search (Smith and Waterman, 1981), faster heuristics solutions were developed, like BLAST (Altschul et al., 1990) or MMseqs (Hauser et al., 2016; Steinegger and Söding, 2017). Having more than one tool to carry out the same task is a mixed blessing. For instance, systematic benchmarks of different MSA algorithms have consistently revealed uneven performances across alternative reference datasets (Santus et al., 2023). A similar situation was reported for read mapping (Donato et al., 2021) or genome assembly methods (Zhang et al., 2022). This situation brings challenges for both users and developers. Users face the complex task of navigating a diverse array of tools, and having to select the most suitable option, considering factors such as accuracy and speed. Meanwhile, developers who investigate and publish novel methods must deploy extensive comparisons to the state-of-the-art.

MSA modeling is a prime example of a problem where the computational complexity has forced pervasive reliance on approximations. MSAs are among the most widely utilized modeling techniques in bioinformatics as they provide the means to compare and analyze genetic sequences, unveiling evolutionary relationships and conserved regions (Van Noorden et al., 2014). This capacity is fundamental for various biological applications, including protein structure prediction, functional annotation, evolutionary modeling, and the understanding of genetic variation (Frazer et al., 2021; Holder and Lewis, 2003; Jumper et al., 2021). The forecast output of large-scale sequencing projects, like the Earth BioGenome Project (Lewin et al., 2022), will present the MSA field with major scale-up challenges requiring improvements well beyond the current state-of-the-art (Santus et al., 2023).

While structure-based approaches to MSA have hitherto been limited by the availability of experimentally determined protein structures, recent progress in protein structure prediction (Abramson et al., 2024; Jumper et al., 2021; Mirdita et al., 2022; Varadi et al., 2022) raises the possibility of utilizing structural information in MSA at a large scale. This additional source of information is especially useful in highly divergent sequences, where the degree of conservation of measured or predicted protein folds can allow more robust alignments than sequence similarity alone (Baltzis et al., 2022b; O’Sullivan et al., 2004), but presents computational and deployment challenges. Recent work on using structurally augmented alphabets to improve the scalability of structural MSA (Gilchrist et al., 2024; van Kempen et al., 2022) confirms the fate of MSA methods as an -omics data integrative procedure. Consequently, a key objective is to provide the community with a generic and flexible deployment platform that integrates and compares all these methods.

Establishing such a unified framework brings several practical challenges. The most obvious issue is that the requirements of the new algorithms can be very diverse. For instance, they may rely on CPU, GPU, or a combination of both, require a local copy or cloud-based access to a large data corpus, and have complex chains of dependencies on libraries and tools. The community has long recognized these problems and it is now considered good practice to deploy scientific computation using workflow management systems (Wratten et al., 2021), offering efficient task orchestration and resource management. Nextflow (Di Tommaso et al., 2017) has become one of the most popular of these solutions (Langer et al., 2024). Nextflow is used by the nf-core community (Ewels et al., 2020), aimed at developing and supporting standardized pipelines to cover all aspects of computational biology. The nf-core ecosystem provides a highly suitable environment for incorporating novel computational developments while ensuring they remain connected to the large corpus of existing methods.

In this application note, we introduce *nf-core/multiplesequencealign*: a framework designed to facilitate MSA deployment while providing rigorous performance evaluation and benchmark reporting. Harnessing Nextflow’s DSL2 extension (Langer et al., 2024), the pipeline’s modular design enables the easy integration of newly developed methods. In this first installment, we have integrated all the methods used for a recent large-scale benchmark (Santus et al., 2023): ClustalO (Sievers et al., 2011), FAMSA (Deorowicz et al., 2016), Kalign3 (Lassmann, 2020), LearnMSA (Becker and Stanke, 2022), MAFFT (Katoh, 2002), MAGUS (Smirnov and Warnow, 2021), Muscle5 (Edgar, 2021), Regressive (Garriga et al., 2019), UPP2 (Park et al., 2022), as well as others widely used tools such as T-Coffee (Notredame et al., 2000) and structure-based aligners: FoldMason (Gilchrist et al., 2024), mTM-Align (Dong et al., 2018) and 3D-Coffee (O’Sullivan et al., 2004). The large-scale capacities of most of these components were previously benchmarked (Santus et al., 2023). Yet *nf-core/multiplesequencealign* is as much a functional pipeline as a seed for an MSA ecosystem in which all members of the community should feel invited to integrate their own MSA methods. Its documentation has therefore been developed for both tool users and developers. This framework should also be considered a pilot for other bioinformatics problems that require the deployment and comparison of alternative modeling methods, such as protein structure prediction, differential expression analysis, and other computationally complex tasks.

## Materials and methods

### Implementation

*nf-core/multiplesequencealign* is written in Nextflow (Di Tommaso et al., 2017) DSL2 syntax (Langer et al., 2024) and is part of the *nf-core* collection of bioinformatics workflows (Ewels et al., 2020). The modular nature of DSL2 extension is crucial to allow 1) the reusability of the individual modules as well as 2) the integration of new components into the pipeline, such as new MSA algorithms or evaluation techniques (see “Integration of new methods”). As required by the nf-core guidelines, each of the modules is provided with a stable software environment either via the conda package manager or software containers such as Docker and Singularity. The implementation in Nextflow combined with the usage of software containers ensures the reproducibility and portability of nf-core/multiplesequencealign across various computing environments (local computers, HPC clusters, and different cloud providers). A contained test dataset is used for continuous integration testing (CI) and provided at https://github.com/nf-core/test-datasets/tree/multiplesequencealign. A full-size dataset is provided at s3://ngi-igenomes/testdata/nf-core/pipelines/multiplesequencealign/1.1.0 and is used to test the pipeline on AWS. The pipeline also comes with extensive documentation of its parameters and usage to be found at https://nf-co.re/multiplesequencealign.

### Input

The minimal input for the pipeline consists of either a FASTA file or a set of protein structures in addition to the specification of which MSA tool to use. The input can be provided in two alternative ways: (1) The parameters *--seqs* and *--pdbs_dir* can be used to provide the FASTA file and/or the set of protein structures respectively, *--tree*, *--args_tree, --aligner,* and *--args_aligner* to provide the directives of which combination of guide tree and aligner to use. This will only allow deploying one dataset using one method for each pipeline run. (2) Alternatively, to deploy the pipeline on multiple samples and multiple tools in parallel, the pipeline’s input must consist of two CSV files. The one received by the parameter *--input* contains the information about the input files, such as the sequences to be aligned, any accessory information to the sequences (e.g., protein structures), and reference alignments to be used for benchmarking. For basic MSA deployment, either the protein sequences or structures are sufficient input, depending on the MSA tools specified. The remaining inputs are optionally used by other modes of the pipeline, such as benchmarking. The second CSV received by the parameter *--tools* contains instructions about which of the tools are to be deployed. Each row specifies one combination of tools, specifically of a guide tree and an assembly method, and the specific combination of flags for both the guide tree and the assembly. A more extensive description of how to build each of the input files can be found in the pipeline’s documentation.

### Integration of new methods

An essential property of *nf-core/multiplesequencealign* is its extensible, modular structure, which allows the incorporation of new methods as they get developed by the community. Adding a new MSA method to the pipeline involves just two steps. First, one needs to provide the tool as a Nextflow module. Ideally, this should be formatted as an nf-core module (https://nf-co.re/modules/) to align with community standards for interoperability and reproducibility. The module’s output is expected to conform to the established standards within its category, e.g., the module of an MSA tool should output an MSA in FASTA format. It is advisable to use existing modules from the same category as templates. These can be found directly in nf-core/modules (https://nf-co.re/modules/) as well as directly installed in the pipeline (https://github.com/nf-core/multiplesequencealign/modules/nf-core). Then, the relevant subworkflow has to be updated to incorporate the new module correctly, using the documentation at https://nf-co.re/multiplesequencealign/docs/usage/adding_a_tool.html and examples from existing tools as a guide. Further instructions for integrating additional modules can be found in the pipeline’s documentation.

## Results

### nf-core/multiplesequencealign deploys and compares various MSA tools: an overview of the pipeline

The pipeline is explicitly designed to deal with protein sequences. As such, it can receive sequences, structures, or a combination of both as input, depending on the requirements of the respective tools (see *Input in Material and Methods*). The *nf-core/multiplesequencealign* pipeline consists of 6 main steps (Figure 1). In the first step, input files are collected, validated, and, optionally, summary statistics are computed (number of sequences to be aligned, sequence length, pairwise sequence similarity). Extra metrics based on structural inputs can also be extracted, such as the pLDDT measure for predicted protein structure quality. The second step involves the computation of the guide tree that may be required by the MSA assembly carried out in step #3. This layout reflects the algorithmic nature of most MSA packages that are based on the progressive alignment algorithm (Hogeweg and Hesper, 1984) in which sequences are first clustered into a binary guide tree and subsequently progressively incorporated into the final MSA following the order indicated by this tree. In *nf-core/multiplesequencealign*, we have explicitly separated steps #2 and #3 to provide a framework that supports the systematic exploration of combinations between guide tree and assembly algorithms. Step #4 is facultative and estimates a consensus model from the computed MSAs using M-Coffee (Wallace et al., 2006). Step #5 evaluates the computed alignments with multiple commonly used quality metrics (see MSA evaluation). The final step (#6) assembles the input statistics and alignment evaluation metrics into a final report. Step #6 optionally also provides the visualization of the rendered MSAs and the corresponding structures using FoldMason (Gilchrist et al., 2024). A key feature of the pipeline, stemming from its Nextflow implementation, is that multiple tools as well as multiple combinations of tools and parameters can be run in parallel. Provided some external quality control is available, such as reference alignments or quality metrics, users can run the pipeline as a benchmark analysis and select the combination of tools most likely to perform well on their production datasets.

**Figure 1:**
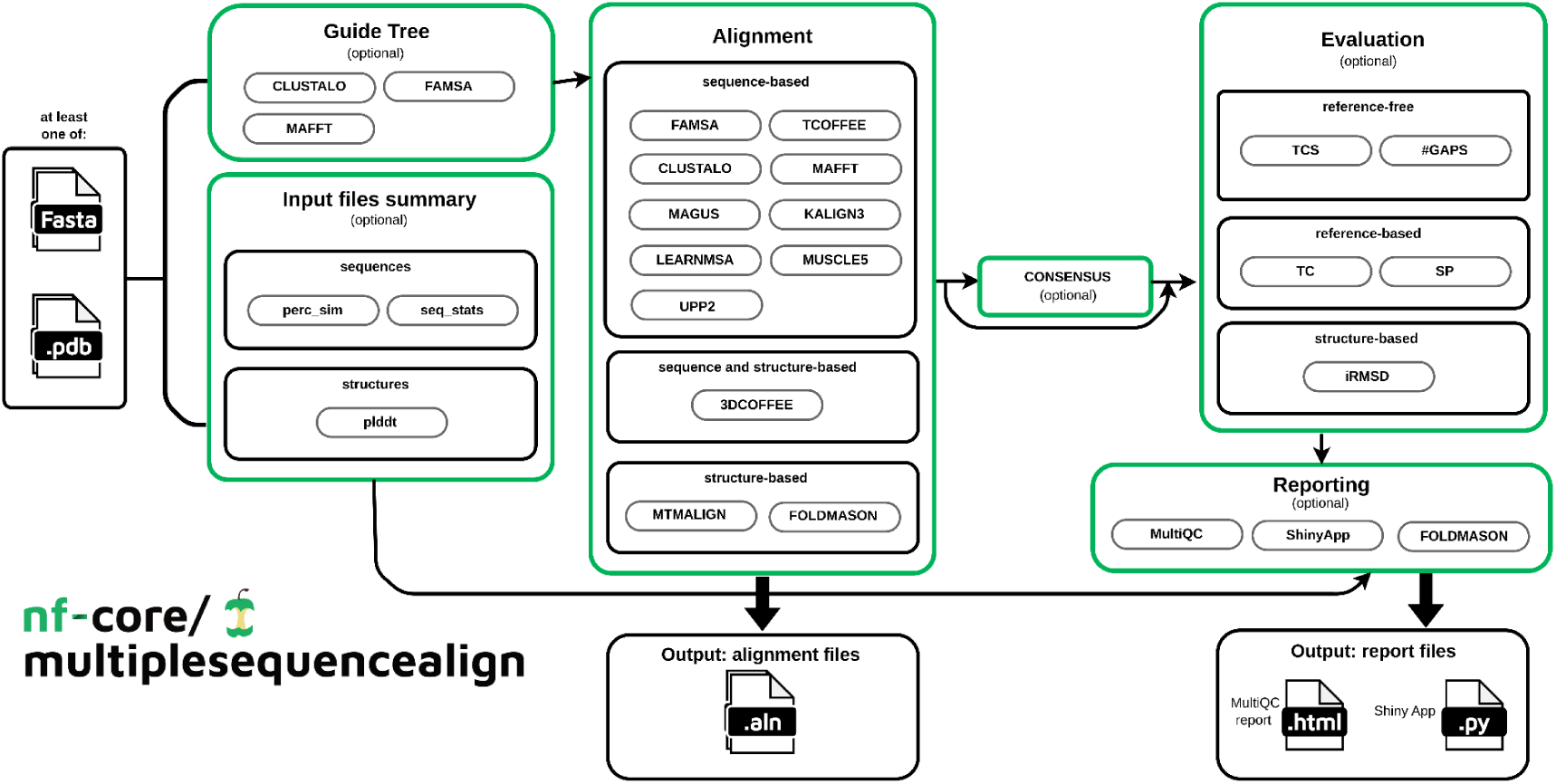
Visual outline of the nf-core/multiplesequencealign pipeline (v1.1.0). First, optionally, the input files are characterized via a collection of summary statistics. Optionally, a guide tree is rendered. Then, the MSA is computed. Finally, various metrics can optionally be calculated to evaluate the tools’ performance. Eventually, these results are summarized in a visual form in MultiQC and a custom shiny app.

This pipeline is therefore a dual tool: (i) a versatile production pipeline allowing the processing of any dataset with a user-defined methodology, and (ii) a benchmark allowing the systematic comparison of alternative MSA procedures and subsequent parameterization of an optimal production pipeline. These two alternative ways of deploying the pipeline are made possible by the toolsheet feature that defines which MSA procedures will be applied to each dataset.

### The toolsheet enables parallel comparison of tools and parameters’ configurations

MSAs, like most bioinformatics tools, rely on complex parameter configurations. Although all possible command line options are part of the same package, in practice, each unique combination of parameters may function as a distinct tool, and sometimes even deploy entirely different algorithms. For instance, in T-Coffee, the default algorithm uses the original consistency-based T-Coffee algorithm (Notredame et al., 2000) while the *-reg* flag triggers a completely different large-scale alignment mode (Garriga et al., 2019). Other popular tools, such as MAFFT, support an even broader range of algorithms (Katoh, 2002). This complexity makes parameterization a feature as important to trace and maintain as the software name and version.

On top of this, each combination of computing elements (e.g., guide tree and assembly) defines a procedure with distinct properties. Each of these procedures is considered separately for benchmarking, as it holds specific properties. The pipeline must account for this complexity, make it traceable, and allow users to unambiguously identify the effect of each component when benchmarking. To guarantee this, in *nf-core/multiplesequencealign* a pipeline benchmarking execution can be specified specified by two input directives: the samplesheet defining the input file (containing the protein sequences, structures, reference alignment, etc.), as used in most nf-core pipelines, and a toolsheet, which declares the tools that will be used and compared. Each line in the toolsheet is considered a distinct procedure in the benchmark and the same tool may appear on several lines with different parameters. Information provided by the toolsheet includes the guide tree method (if applicable), the assembly method, and the command line parameters. Each combination of tools defined in the toolsheet is then computed for each sample specified in the samplesheet. This process can be seen as a Cartesian product of the sample and toolsheet. This results in a collection of MSAs ready to be compared with one another or with reference datasets. In a production use case where a single output is desired, the possibility exists to limit the toolsheet to a single tool only or to use the *--tree*, *--args_tree*, *--aligner*, *--args_aligner* parameters (see Methods).

### Guide tree computation

Guide trees, which are a hierarchical clustering of input sequences, define the order in which sequences are aligned. They are a prerequisite for most progressive MSA methods (Santus et al., 2023). Guide trees have been shown to have a major impact on MSA accuracy (Garriga et al., 2019). Their computation can be a bottleneck in large datasets as they typically rely on a distance matrix of all sequences to be calculated, requiring O(N^2^) operations. As of version 1.1.0, the pipeline includes FAMSA (Deorowicz et al., 2016) to support the stand-alone computation of the most common types of guide tree methods, specifically Neighbor-joining (Saitou and Nei, 1987), the unweighted pair group method with arithmetic mean (UPGMA) (Sokal and Michener, 1958), single linkage (SL) and PartTree (Katoh and Toh, 2007) using either SL or UPGMA. Multiple flavors of the PartTree heuristic can be also deployed using MAFFT (Katoh and Toh, 2007), reported in detail in Table 2. The pipeline also supports the ClustalO (Sievers and Higgins, 2014) implementation of mBed trees, which are based on a combination of k-means clustering and sequence embeddings (Blackshields et al., 2010). The guide tree computation step has been designed modularly to allow the easy incorporation of new guide tree methods users may be interested in.

### MSA assembly

Assembly refers to the step where the final MSA is constructed by combining all the sequences, a procedure common to all MSA tools, regardless of their dependence on a guide tree (Chatzou et al., 2016; Santus et al., 2023). The pipeline integrates some of the most widely used large-scale assembly methods (listed in Table 1), and like the other sections of the pipeline, it has been specifically designed in a modular way to support the integration of novel tools as they get developed (see *Integration of new methods)*.

**Table 1:**
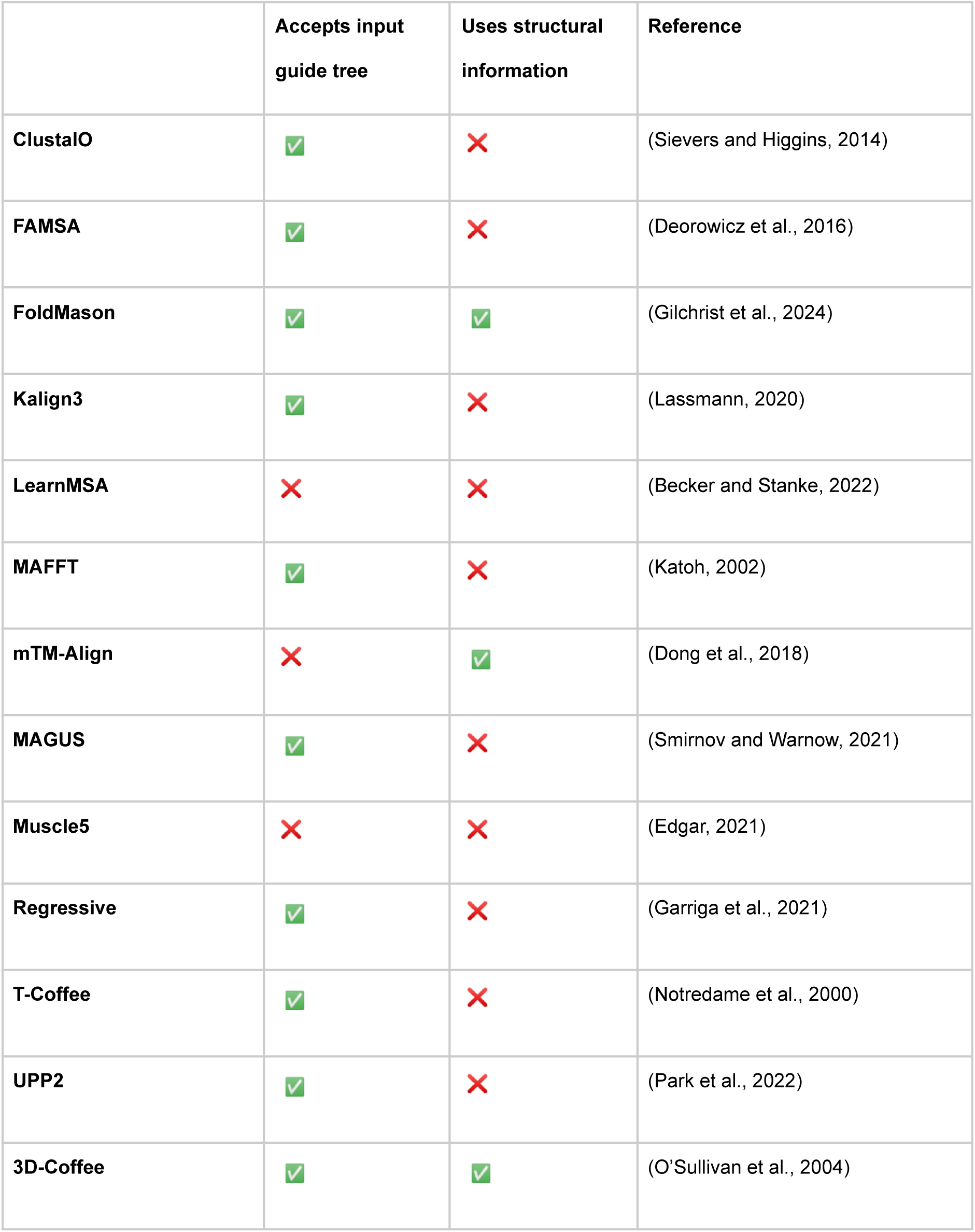
MSA Methods included in nf-core/multiplesequencealign v1.1.0. The column “Explicit guide tree definition” specifies whether the tool can accept an explicit definition of a guide tree to be used. Some tools may use one internally but do not accept the input of a custom guide tree. The column “Uses structural information” specifies whether the aligner is structural.

|  | Accepts input<br>guide tree | Uses structural<br>information | Reference |
| --- | --- | --- | --- |
| **ClustalO** | ✓ | ✗ | (Sievers and Higgins, 2014) |
| **FAMSA** | ✓ | ✗ | (Deorowicz et al., 2016) |
| **FoldMason** | ✓ | ✓ | (Gilchrist et al., 2024) |
| **Kalign3** | ✓ | ✗ | (Lassmann, 2020) |
| **LearnMSA** | ✗ | ✗ | (Becker and Stanke, 2022) |
| **MAFFT** | ✓ | ✗ | (Katoh, 2002) |
| **mTM-Align** | ✗ | ✓ | (Dong et al., 2018) |
| **MAGUS** | ✓ | ✗ | (Smirnov and Warnow, 2021) |
| **Muscle5** | ✗ | ✗ | (Edgar, 2021) |
| **Regressive** | ✓ | ✗ | (Garriga et al., 2021) |
| **T-Coffee** | ✓ | ✗ | (Notredame et al., 2000) |
| **UPP2** | ✓ | ✗ | (Park et al., 2022) |
| **3D-Coffee** | ✓ | ✓ | (O'Sullivan et al., 2004) |

**Table 2:** Guide tree rendering methods included in nf-core/multiplesequencealign v1.1.0. The “Guide tree” column specifies the guide tree and its associated command line parameters. The “Distance measure”, “Core algorithm”, and “Speed-up heuristic” columns define the distance measure, core algorithm, and specific heuristic used when the guide tree specified in the “Guide tree” column is selected.

| Guide tree | Distance measure | Core algorithm | Speed-up heuristic |
| --- | --- | --- | --- |
| MAFFT | k-mer-based | UPGMA + single linkage combined |  |
| MAFFT --minimumlinkage | k-mer-based | single linkage |  |
| MAFFT --averagelinkage | k-mer-based | UPGMA |  |
| MAFFT --parttree | k-mer-based | single linkage + UPGMA combined | PartTree |
| MAFFT --dpparttree | dynamic programming alignment-based | single linkage + UPGMA combined | PartTree |
| MAFFT --fastaparttree | FASTA alignment-based | single linkage + UPGMA combined | PartTree |
| CLUSTALO | sequence embedding + approx. alignment | UPGMA | bisecting K-means |
| FAMSA | longest common subsequence-based | single linkage |  |
| FAMSA -gt upgma | longest common subsequence-based | UPGMA |  |
| FAMSA -gt nj | longest common subsequence-based | neighbour joining |  |
| FAMSA -parttree | longest common subsequence-based | single linkage | PartTree |
| FAMSA -gt upgma -parttree | longest common subsequence-based | UPGMA | PartTree |
| FAMSA -medoidtree | longest common subsequence-based | single linkage | MedoidTree |
| FAMSA -gt upgma -medoidtree | longest common subsequence-based | UPGMA | MedoidTree |

In the current implementation, iterative algorithms occasionally employed for MSA computation are carried out within the framework of their dedicated software, even when these iterations involve re-estimating a guide tree (Liu and Warnow, 2014; Sievers et al., 2011; Sievers and Higgins, 2014).

### Generation of a consensus MSA

Dealing with several alternative models estimated on the same data presents a significant challenge, especially when no single objective measure is available to unequivocally decide which model should be preferred over the others. Consensus MSAs provide a convenient way to address this issue as they allow the information content of several MSAs to be merged into a single model. This integration is especially useful for identifying highly supported regions in multiple MSAs. These have been shown to match the structural similarity of proteins more closely than less supported regions (Wallace, 2006). The current version of the pipeline supports the generation of consensus alignments using the M-Coffee protocol. Under its formulation, all pairwise projections are extracted from the initial MSAs and used to populate a position-specific scoring scheme based on consistency. This scheme is then used to estimate a progressive MSA. This MSA constitutes an empirical consensus that has been shown to have a high average sum-of-pairs score, comparable to the scores of the individual MSAs. In other words, the consensus MSA effectively captures the shared characteristics of the constituent alignments while preserving a level of quality similar to that of the original inputs. The consensus MSA also comes with several global and local metrics, allowing an estimate of the global agreement between the alternative alignments.

### MSA evaluation

A significant hurdle in the development of novel MSA algorithms lies in the intricacy of their evaluation. For a long time, the lack of references resulted in the MSA problem being defined in terms of string matching. Under this formulation, an MSA is optimal if it minimizes the alignment score between the projected alignment of every pair of sequences. This formulation is known as the Sum of Pairs (SoP). In practice, the SoP formulation constitutes, more or less explicitly, the objective function of most multiple sequence alignment algorithms. The SoP is, however, known to be limited in its capacity to produce structurally optimal (Notredame, 1996) and biologically meaningful MSAs. The question of how to evaluate MSA accuracy therefore remains partly open, especially considering the lack of an absolute, formal definition of MSA correctness. Indeed, the final evaluation in practice will often depend on domain-specific criteria that may arbitrarily incorporate elements of evolutionary, structural, or functional perspectives on correctness.

In biological terms, an MSA may be described as the product of an evolutionary process and should as such precisely reflect the unique evolutionary history of the observed sequences. To be of any use, this formulation requires prior knowledge of the correct underlying phylogeny, an information usually not available. However, evolutionary correctness is only one perspective when evaluating MSAs.

Often, MSAs are used to model structural similarity among homologous sequences (Jakočiūnas et al., 2018), in which case correctness is defined as a geometric problem. While structure- and evolution-based criteria for evaluating MSAs tend to have some level of agreement, their equivalence remains disputed (Chang et al., 2014; Liu and Warnow, 2014; Löytynoja, 2014). In practice, the most commonly used validation method has been the comparison of sequence-based MSAs with reference structure-based counterparts. These comparisons can either be carried out using the SoP between the two MSAs or the comparison of entire columns (Total Column, TC). However, reference datasets of structure-based MSAs are not entirely straightforward to establish - the computation of structure-based MSAs is also an NP-Hard problem. This extra hurdle has resulted in the establishment of several alternative reference MSA collections (Bahr, 2001; Edgar, 2004; Mizuguchi et al., 1998). Alternative alignment-free methods were also reported, based on the comparison of structural properties (F. Armougom et al., 2006; Gilchrist et al., 2024; Holm, 2020; Mariani et al., 2013; Taylor, 2000; Zhang and Skolnick, 2004).

The current implementation of the pipeline features the possibility to do an SoP and a TC comparison between the MSA produced by any procedure on any input sample and a user-provided sample-specific reference (specified in the “reference” column of the samplesheet), using these metrics as implemented in the T-Coffee package (Notredame et al., 2000). The pipeline also supports two alignment-free metrics, the TCS score that estimates the agreement between an MSA and the optimal alignments associated with its pairwise projections (Chang et al., 2014), and the normalized intra-molecular root mean square deviation (NiRMSD), if at least two protein structures are provided for the input sample (Fabrice Armougom et al., 2006). The NiRMSD quantifies the structural agreement of the aligned sequences. The pipeline can also provide simple summary metrics, including the count of gaps in the alignment (Table 3).

**Table 3:** Evaluation methods available in nf-core/multiplesequencealign v1.1.0. The column “reference-based” specifies whether the tool needs a reference alignment to compute the tool’s performance. The column “structure-based” specifies whether structures are needed for the evaluation. All metrics, except for the number of gaps, are implemented in the T-Coffee/alncompare module.

| Method | Reference-based | Structure-based | Reference |
| --- | --- | --- | --- |
| Sum-of-Pairs (SP) | ✓ | ✗ | (Bahr, 2001; Notredame, 1996) |
| Total column (TC) | ✓ | ✗ | (Bahr, 2001) |
| TCS | ✗ | ✗ | (Magis et al., 2014) |
| Number of Gaps | ✗ | ✗ | Custom script |
| NiRMSD | ✗ | ✓ | (Fabrice Armougom et al., 2006) |

We expect the effective evaluation of MSAs to remain a very active field of investigation, especially under the pressure of deep-learning-based methods, now routinely used to scan MSAs for pathogenic mutations (Frazer et al., 2021) and the development of new structural aligners in light of the wide availability of predicted protein structures (Gilchrist et al., 2024). For this reason, the implementation of the evaluation section has also been explicitly designed to facilitate the integration of novel evaluation metrics.

As datasets grow, feasible resource requirements are crucial to maintaining the practicality of these tools. Resource utilization, including CPU usage, memory consumption, and runtime, is therefore a crucial factor in tool evaluation. To address these needs, nf-core/multiplesequencealign leverages Nextflow’s native tracing and the nf-co2footprint plugin to provide a final report that includes resource utilization metrics and estimated CO2 emissions.

### Summary Report

The pipeline offers an optional summary report detailing the metrics calculated on the input datasets and the evaluation of the resulting alignments using MultiQC (Ewels et al., 2016), (Figure 2). Additionally, if structural data is provided, Foldmason (Gilchrist et al., 2024) enables user-friendly visualization of the rendered MSAs along with their corresponding structures. *nfcore/multiplesequencealign* also provides a shiny-based browser interface to explore the results, as illustrated in Figure 3. A standardized way to systematically evaluate the impact of certain input features – captured by the summary statistics reported by *nfcore/multiplesequencealign* – can provide MSA tool developers with a deeper understanding of when MSA tools perform well, and when they fail. This can help optimize tools for specific situations or even inspire the development of new tools exploiting different kinds of signals in the data.

**Figure 2:**
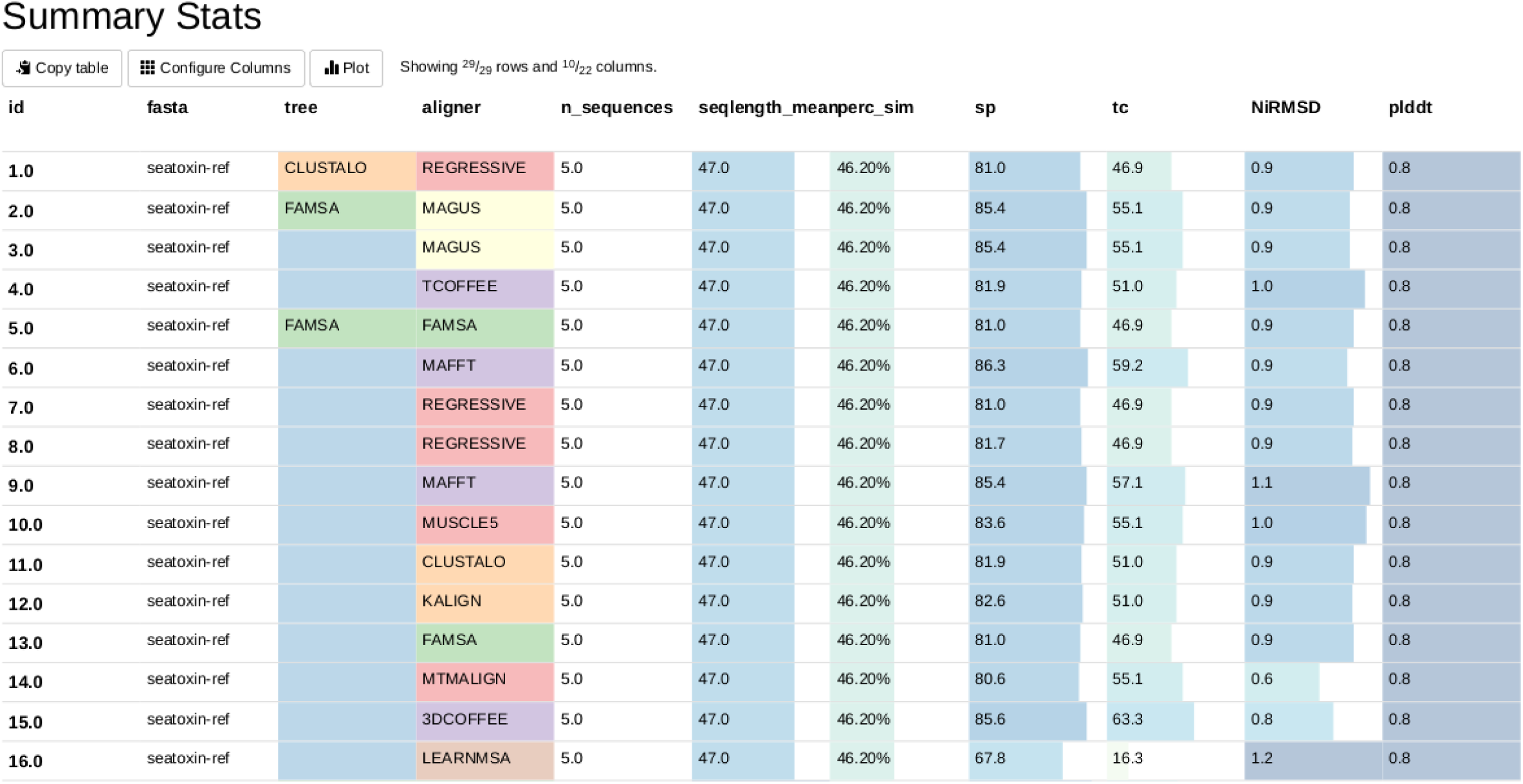
MultiQC summary table. A single dataset (column fasta) was used to test a variety of tools (combination of columns tree and aligner). The remaining columns show summary statistics that characterize the input files (n_sequences, seqlength_mean, perc_sim) and the tool performance evaluation (sp, tc, NiRMSD, plddt).

**Figure 3:**
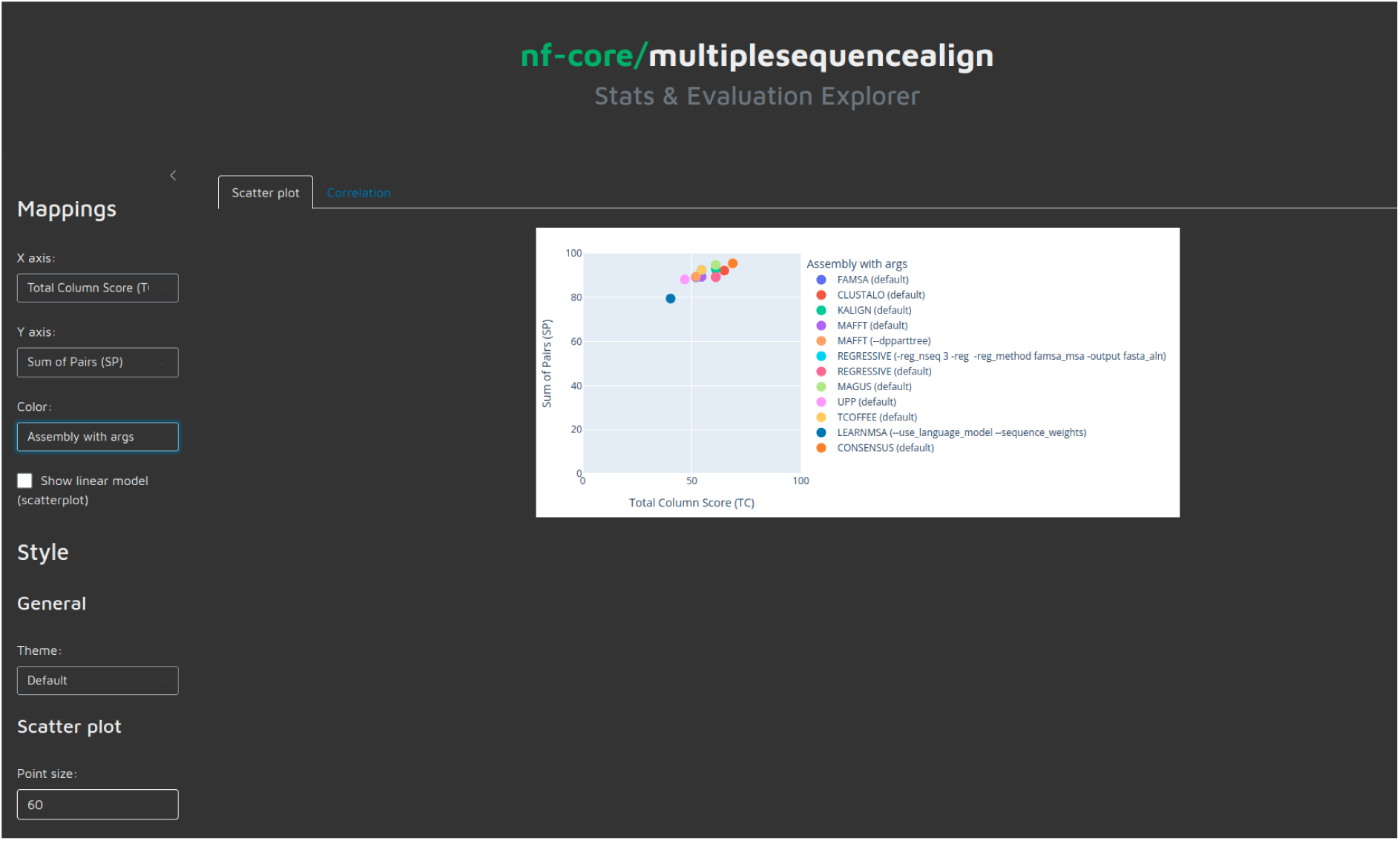
Screenshot of the shiny app of the pipeline. A shiny app is created to explore the results of the analysis. All metrics (summary statistics about the input files, type of tools and parameters deployed, evaluation metrics, time, and memory requirements) are available for interactive exploration.

### Pipeline chaining

A key design principle within the nf-core community indicates that each pipeline should focus on a particular type of analysis. For instance, pipelines may be dedicated to RNA-Seq (nf-core/rnaseq), variant calling (nf-core/sarek), (Hanssen et al., 2024), or protein folding (nf-core/proteinfold) (Baltzis et al., 2022a). However, it is extremely common for the output of one pipeline to serve as the input for another. For instance, the *nf-core/proteinfold* pipeline accepts one or more FASTA files and utilizes various protein structure prediction tools, such as AlphaFold2 or ColabFold, to generate predicted structures. The obtained protein structures can be an input for the *nf-core/multiplesequencealign* workflow. Therefore, integrating the results of *nf-core/proteinfold* into *nf-core/multiplesequencealign* is bound to be a common use case. *nf-core/multiplesequencealign* supports this integration, as detailed in the documentation.

## Discussion

Despite its central role in biological analysis, the effective deployment of MSA computation remains hampered by rapidly growing genomics datasets, the high computational complexity of available algorithms, and the lack of deployment and documentation standards. On top of this, the wide range of downstream uses of MSAs, spanning from orthology detection and functional genomics to phylogeny reconstruction and protein structure prediction, each brings its own challenges and requirements. Combined with the lack of clear, unequivocal solutions, this has resulted in an extremely fragmented software landscape. The sequence alignment Wikipedia page lists 52 distinct MSA packages [https://en.wikipedia.org/wiki/List_of_sequence_alignment_software#Multiple_sequence_alignment] developed over more than 35 years. The packaging, usage, and output formats of this software are not standardized in any way and remain extremely diverse. This makes general usage and thorough comparisons difficult: except for a recent review on large-scale alignments this work is based upon, we are not aware of any comparative study including more than 10 MSA packages. Yet, the most ancient of these solutions were developed at a time when pressure on hardware, especially concerning memory, was stronger than it is today. It cannot be ruled out that older algorithms may contain the seed of future solutions to the current scale-up problem. Maintaining the memory of the field is, therefore, an issue of prime practical interest. Achieving this goal will require the community to come together to develop an ecosystem in which all the relevant methods can be dynamically maintained and systematically compared. We hope to lay the groundwork for such an ecosystem with this pipeline.

*nf-core/multiplesequencealign* paves the way toward the establishment of an exhaustive and possibly automatically updated reference benchmark workflow for MSA tools. The requirement of live standardized benchmarks is crucial in the case of complex computational processes that combine various heuristics. Often enough, these algorithms are iteratively improved by their authors. Yet, whenever an updated algorithm is released, it is down to the data analysis team to establish whether improvements carried out independently across combined algorithms are additive, synergistic, or antagonistic. It is also down to the users to determine a suitable trade-off between the energetic cost of their computation and the resulting added value. In practice, one may expect this assessment to be domain-dependent or even test-case-dependent. Facilitating benchmarking is therefore a major goal for the long-term sustainability of computational biology. We expect that the establishment of live benchmarkings will be needed well beyond MSA, across all domains relying on fast-evolving algorithms.

While our framework doesn’t directly address the challenges associated with the lack of clear and unbiased benchmarking datasets for MSAs, it offers an approach that is both dataset-agnostic and inclusive of multiple evaluation methods deemed of interest by different communities of users. This is particularly important in light of the myriad downstream uses of MSAs that have led to the development of a large number of accuracy metrics over the years. Establishing the relative merits of these alternative evaluations has been proven difficult and it is common practice to report readouts on two or more reference datasets or stand-alone metrics. Once populated with all the relevant evaluation methods, our pipeline should allow for more systematic comparisons. More importantly, it will allow users to identify which metrics best reflect their needs and choose their alignment strategy accordingly. Since the metric defines the biological optimality of an MSA, its selection is as important as the algorithmic optimization capacities. Yet these two levels of optimization - the computational and the biological - have so far remained largely disconnected. Here, we present the community with a standardized, documented, and extensible system in which the dependencies between optimization and evaluation can be explored.

Our pipeline also hints in the direction of template-based sequence alignments, a concept pioneered in 3D-Coffee nearly two decades ago. Template-based alignments rely on the concept that the alignment of a protein sequence may be enhanced by adding an additional layer of information. The template is a re-encoding of the sequence in which specific features are collected that lend themselves to the establishment of an alignment without using the original sequence information. Structural data is the most commonly used kind of information that can be associated with sequence data, which the pipeline can already use as a template. Over the years, other types of templates have been developed such as RNA secondary structure-based (Wilm et al., 2008) and evolutionary (Chang et al., 2012) templates. In theory, any data that can be mapped onto DNA, RNA, or protein sequences is suitable to become a template. For instance, the notion of functional templates using functional genomic information associated with sequences remains to be explored. It could provide important developments in DNA analysis, thanks to the wide availability of DNA functional data collected by projects like ENCODE or via prediction tools (Avsec et al., 2021). The consistency-based approach of T-Coffee in which templates were originally developed is especially suitable for this kind of integration. We anticipate the use of additional information in MSA will become standard practice, especially for the alignment of regulatory regions, turning MSA into a multi-omics problem. We expect an extension of the pipeline to allow for these analyses to be an important future milestone.

The maintenance of a common hub for MSA methods is a long-term commitment. Because of the inherent computational complexity of the problem, MSA methods are bound to keep evolving, following the information content, scale, and downstream applications of sequences generated by biologists. By providing a common roof to a large number of methods, we expect that this pipeline will allow the gradual deconstruction of existing methods into smaller, modular algorithmic components. Indeed, and as reflected in the current layout of the pipeline, MSA algorithms consist of multiple, independent steps. Yet, historically, most packages have been distributed as self-contained units. This approach has often led to the reimplementation of existing components, tightly integrated at the core of each package. In the case of MSAs, some sub-procedures may be for instance the computation of the distances used for some guide tree computation, the guide tree computation itself, and the assembly. At times, this process makes it difficult to identify which component drives performance improvement. Addressing this problem will require systematically deconstructing every method into a collection of modules, each implementing a step of the original algorithm. The basic principle of Nextflow and their implementation in *nf-core/multiplesequencealign* provide an ideal framework for such deconstruction. This process requires a wide effort by the method development community, yet the modular nature of Nextflow allows it to be done in a progressive and decentralized manner by leveraging the power and resources provided by the nf-core community and its many members.

## Data availability

nf-core/multiplesequencealign is available on GitHub under the nf-core organization (https://github.com/nf-core/multiplesequencealign). It is released under the MIT license. Test datasets are available at https://github.com/nf-core/test-datasets/tree/multiplesequencealign and s3://ngi-igenomes/testdata/nf-core/pipelines/multiplesequencealign/1.1.0.

## Acknowledgments

We would also like to thank the nf-core community for the continuous support during the development of the pipeline. A full list of nf-core community members is available at https://nf-co.re/community.

## Author contributions

L.S., J.E.C. and C.N. designed the project. L.S. led the implementation of the project. J.E.C., L.R., J.M., A.V., I.T., L.M. contributed to the implementation. A.B., E.W.F., P.D.T., E.G., A.G., S.D., C.G. and M.S. contributed ideas to the initial design of the project. All authors wrote the manuscript.

